# Genus-wide re-speciation of *Corynebacterium* and genomic characterisation of 87 sinus isolates resolves taxonomic conflicts and identifies four candidate novel species

**DOI:** 10.64898/2026.10.13.758830

**Authors:** Isabella Burdon, James Peter Psaltis, Nicholas Christopher Vrodos, Jeffrey Sung, Lindsay Durr, Emma Barry, Clare Cooksley, Sarah Vreugde, Peter John Wormald, Alkis James Psaltis

## Abstract

*Corynebacterium* species are common members of the sinonasal microbiome and have shown conflicting associations with sinus health and chronic rhinosinusitis (CRS) in genus-level surveys. To better resolve the taxonomy and species-level functions of sinus corynebacteria, we generated complete long-read genome assemblies for 87 isolates, sourced from the middle meatus of 71 patients (51 with CRS, 20 healthy controls). We re-derived species boundaries de novo across the whole genus by average nucleotide identity (ANI), clustering 4,661 quality-filtered genomes retrieved from public databases and delineating higher-order species complexes. Our re-speciation assigned a species to 282 unlabelled corynebacteria, and corrected taxonomy for 266 NCBI accessions. We identified four novel candidate species from our isolate collection, and 39 public genomes fell within these species’ boundaries. Whole-genome assignment reclassified 41 of our 87 isolates relative to mass-spectrometry (MALDI-TOF MS) identification, which only resolved to the species-complex level. Comparison of genome annotation characterised accessory functions that partitioned along species-complex lines, including glycogen and glycan utilisation, fatty-acid biosynthesis, cofactor (biotin and thiamine) auxotrophy, and iron-acquisition systems. Virulence and antimicrobial-resistance determinants were sparse, and the diphtheria toxin gene was universally absent. Acquired antimicrobial resistance was enriched on plasmids, whilst virulence genes were depleted from horizontally transferred compartments. Contrary to its frequent report in sinus microbiome studies, no *C. tuberculostearicum* genome publicly accessible via NCBI was found to be isolated from the sinonasal tract. Continued genome-level interrogation of *Corynebacterium* and contribution of sinus-commensals to reference databases will improve species-level understanding of the role of corynebacteria in the sinonasal microbiome.

**Highlights:**

- Type-strain-anchored re-speciation of corynebacteria reassigns hundreds of genomes
- Survey and deposition of 87 *Corynebacterium* genomes, and 4 candidate novel species
- *C. nasorum* not *C. tuberculostearicum* is the sinonasal member of its species complex
- Species diverge in glycan, lipid, cofactor and iron-acquisition strategies
- Virulence is sparse; antimicrobial resistance is also rare but enriched on plasmids

## 1. Introduction

*Corynebacterium* species are salient, yet poorly characterised members of the human microbiome. This genus spans major human pathogens including *Corynebacterium diphtheriae*, and *Corynebacterium striatum* to proposed commensal species such as *Corynebacterium accolens*. In the sinuses, the most commonly referenced corynebacteria include *C. accolens, C. tuberculostearicum, C. pseudodiphtheriticum* and *C. propinquum* (Tran et al., 2024). Study of the sinus microbiome has garnered interest as a means to deepen our understanding of chronic rhinosinusitis (CRS) aetiopathogenesis and recalcitrance.

While some sinus microbiome studies link the presence of corynebacteria to health (Huffines et al., 2024; Kim et al., 2020; Paramasivan et al., 2020; Zhu et al., 2023), others find no significant difference in patient outcomes (Gan et al., 2021; Lux et al., 2020; Rom et al., 2019), or even find members of the *Corynebacterium* genus to be pathobionts associated with CRS (Abreu et al., 2012; De Boeck et al., 2019). This speaks to the challenge of drawing conclusions from microbiome studies that only resolve to the genus level. Genus-level aggregation both limits the clinical applicability of such studies, and fails to capture the ecologically relevant interactions that occur between species (Bassiouni et al., 2020; Huffines et al., 2024).

In studies of human-associated microbiota, corynebacteria have demonstrated highly species-specific relationships with co-colonising bacteria. For example, *C. accolens* and *C. pseudodiphtheriticum* antagonise the growth of *Staphylococcus aureus* (Escapa et al., 2018; Hardy et al., 2019; Huffines et al., 2024; Menberu et al., 2021; Yan et al., 2013), whilst *Streptococcus pneumoniae* growth is also restricted in the presence of *C. accolens* (Bomar et al., 2016). Understanding these interactions between species is paramount to our ability to distinguish between commensal and pathogen in the sinus microbiome (Escapa et al., 2018).

In-depth interrogation of *Corynebacterium* species through genome assembly and annotation is a foundational step towards resolving this area of contention (Tran et al., 2024). Compared with traditional culture- and molecular-based methods, genome-wide studies provide greater insight into bacterial taxonomy and function (“A genomic approach to microbiology,” 2019; Goyal et al., 2023). Moreover, generation of complete bacterial genomes facilitates expansion of reference databases, thereby supporting downstream genomic and microbiome analyses (Kim et al., 2022).

The *Corynebacterium* genus is under active revision. In 2025, *Corynebacterium rhinophilum* was characterised and split from its neighbour *C. accolens* (Popowitch et al., 2025). Further still, the species *Corynebacterium nasorum* and *Corynebacterium hallux* have recently been identified as novel members of the *C. tuberculostearicum* complex (E. Popowitch et al., 2024). This shifting landscape raises an issue for the fidelity of metagenomic microbiome study, which is particularly contingent upon the phylogenetic accuracy, extent of representation, and quality of genomes in the selected reference database (Smith et al., 2022).

To this end, databases such as the Genome Taxonomy Database (GTDB) have been created to standardise taxonomy, remove duplicate and low-quality genome assemblies and provide regular updates (Parks et al., 2022). However, in capturing phylogenetic breadth, the GTDB introduces species and genus epithets that clinical microbiology does not use. Using the same methods of accurate taxonomic assignment but restricting to names approved by the International Code of Nomenclature of Prokaryotes (ICNP) could contribute towards more clinically translatable metagenome studies.

This study aims to re-derive *Corynebacterium* species boundaries, assigning genomes under validly published names, and placing 87 complete long-read *Corynebacterium* genomes from our sinus biobank within this framework. This is implemented by an open-source pipeline which provides a reproducible, type-strain-anchored framework for genus-wide re-speciation that can be updated as public databases change. Further, we aim to evaluate encoded differences in *Corynebacterium* genomes at the species level to generate mechanistic hypotheses.

## 2. Materials and Methods

### Collection of isolates, culture condition and identification

Ethics approval for clinical isolate collection was provided by the Central Adelaide Local Health Network Human Research Ethics Committee (HREC 10604). Guarded Σ-Transwabs (Medical Wire & Equipment, Corsham, United Kingdom) were taken from the middle meatus of patients under endoscopic guidance prior to surgery and stored in liquid amies at 4°C for transport. 50µL of swab eluate was streaked across TSA plates supplemented with 0.5% TWEEN-80 (TSA-T80) (Sigma Aldrich, Bayswater, Australia) and plates were incubated at 37°C. For each unique morphology, single colonies were selected and purified by sequential sub-culture, followed by identification using spectra acquired by ultrafleXtreme and ultraflex III MALDI-TOF MS mass spectrometers (Bruker, Preston, Australia) (University of Adelaide Proteomics Centre, Adelaide, Australia). Purified isolates were stored at −80°C in 1mL of tryptone soya broth (TSB) (OXOID, Vic, Australia), supplemented with 0.5% TWEEN-80 (TSB-T80) and 20% glycerol (Sigma Aldrich).

### Preparation of bacteria and DNA extraction

Liquid cultures were prepared by suspending colonies grown on TSA-T80 plates in TSB-T80 and incubating overnight at 37°C with 180rpm shaking. Bacteria were pelleted (4000rcf, 5mins), followed by DNA extraction with the PureLink™ Microbiome DNA Purification Kit (Thermo Fisher, Thebarton, Australia) with minor adjustments. Briefly, bacterial lysate was prepared by resuspending the bacterial pellet in 800µL of lysis buffer and 100µL of lysis enhancer, then bead beating 4mins at 30Hz using a TissueLyser II (Qiagen, Clayton, Australia). DNA was then bound to the spin column and washed with two wash steps before elution. DNA was assessed for quality and quantity by both Nanodrop spectrophotometer (Thermo Fisher) and Qubit 3.0 Fluorometer (Thermo Fisher) using the high sensitivity kit (Thermo Fisher) per the manufacturer’s instructions.

### Library preparation and sequencing

Libraries were prepared for long read whole genome sequencing using the Oxford Nanopore Rapid Sequencing Kit V14 (SQK-RBK114.96) (Oxford Nanopore Technologies, Oxford, United Kingdom) as per the manufacturer’s protocol with an additional AMPure bead clean up step prior to pooling to prevent barcode bleed. Briefly, 11.5µL of AXP beads were added to each barcoded sample. Samples were pelleted using a magnetic rack, the supernatant was discarded and the pellet was washed twice with 500µL 80% ETOH. 11.5µL elution buffer was added, and samples were incubated for 10mins at room temperature with end over end rotation. Finally, beads were pelleted and 11µL of eluate was retained from each sample and pooled together before being concentrated using AMPure beads per the original protocol. Rapid adapter was added and the library was then loaded into a PromethION flow cell in the P2 Solo (Oxford Nanopore Technologies). Raw signal data was base-called using Dorado’s super-accurate model (Oxford Nanopore Technologies).

### TAXANALYSER bacterial genomes pipeline

A bioinformatics pipeline was established to handle computational analyses. This software “analyse bacterial genomes” is published as part of the open-source TAXANALYSER suite (https://gitlab.com/isabella-burdon/taxanalyser; Apache-2.0). In the present context, this pipeline handled all stages from concatenation and quality filtering of raw FastQ files, assembly (Hybracter), genome QC (CheckM2) and deduplication, GTDB-Tk genus confirmation, annotation (Bakta/Baktfold, KofamScan, dbCAN, antiSMASH, AMRFinderPlus, RGI, VFDB, eggNOG-mapper, mobilome). Concurrently the pipeline facilitated genus-wide NCBI genome and metadata fetch, LPSN-anchored re-speciation (skani/skDER), and per-species pangenomes (PPanGGOLiN). These pipeline stages are detailed below.

### Genome assembly, quality assessment and deduplication

For each barcode, raw FastQ files were concatenated and files were cleaned to remove reads with an average Phred score less than 10 or length less than 1000bp. Genomes were assembled with Hybracter [v0.11.1] using an estimated genome length of 2.2Mbp and a predicted genus of *Corynebacterium* (Bouras et al., 2024). The completeness and contamination of these genomes were assessed with CheckM2; genomes that passed this stage had scored a completeness ≥95% and contamination ≤5% (Chklovski et al., 2023; The Genome Standards Consortium et al., 2017). Next duplicate genomes were identified; these were defined as assemblies that shared an average nucleotide identity (ANI) ≥99.9% as calculated by skani, and with <1000 SNPs and <75 InDels as enumerated by the NUCmer/dnadiff modules of MUMmer4 (Marçais et al., 2018; Shaw and Yu, 2023; Zhang et al., 2024). Duplicate genomes from the same patients were reduced to the single best assembly and dropped from the dataset. Of the 94 isolates sequenced, 73 assembled into complete chromosomes, and 71 were retained after deduplication to be included in the present study.

### Taxonomic assignment and novel species lineages

Genus level taxonomy of clinical isolates was confirmed with the genome taxonomy database tool kit [GTDB-Tk v2] (Chaumeil et al., 2022). To define the within-genus taxonomic framework, we re-derived species boundaries de novo across the *Corynebacterium* genus. A curated set of reference genomes was first assembled, comprising validly published *Corynebacterium* species type strains according to the List of Prokaryotic names with Standing in Nomenclature [LPSN], together with one representative for each GTDB genomospecies cluster not encompassed by LPSN (Parks et al., 2022; Parte et al., 2020). Two species in pre-print with valid type strains were supplemented to this list: *Corynebacterium nasorum* (type strain KPL3804 = ATCC TSD-439 = DSM 117767) and *Corynebacterium hallux* (type strain CTNIH22 = ATCC TSD-435 = DSM 117774) (E. B. Popowitch et al., 2024). The entire *Corynebacterium* genus was then downloaded with NCBI Datasets [5,055 distinct assemblies; accessed 8 August 2026] and pooled together with the quality-filtered, de-duplicated clinical isolates (O’Leary et al., 2024). Metadata for the genus-wide reference set was extracted alongside the genomes. Assembly level, host, country (from geo_loc_name) and isolation source were taken from the NCBI Datasets/BioSample metadata; free-text host and body-site values were mapped to controlled categories by keyword matching with manual curation, and genomes lacking a field were excluded from that field’s denominator.

Reference genomes were retained if they passed a locally computed CheckM2 filter of ≥95% completeness and ≤5% contamination, with type strain genomes excepted from this filter. This yielded 4,661 genomes which were pooled for all-versus-all ANI. All-versus-all ANI was calculated across the pool with skani and species were partitioned de novo with skDER in greedy mode at a 95% ANI and 50% alignment-fraction (AF) boundary (Chun et al., 2018; Jain et al., 2018; Parks et al., 2020; Salamzade et al., 2025; Shaw and Yu, 2023). GTDB assignments were retained as a comparison layer and never defined a species boundary. Each species cluster was classified and named by its composition. If a cluster contained a validly published type strain it was designated that name (Supplementary Figure 1). Type-less clusters were labelled by their nearest type strain within 90% ANI and defined as candidate novel species. These candidate novel species were confirmed where digital DNA-DNA hybridisation (dDDH) to the nearest type strain was <70% [formula d4, GGDC/TYGS] (Meier-Kolthoff et al., 2013; Meier-Kolthoff and Göker, 2019). If a cluster contained no type strain and at least one clinical isolate, the most ANI-central clinical isolate was designated as its proposed type strain. The skDER species clusters were confirmed via two methods; firstly, by their agreement (adjusted Rand index) with an independent average-linkage clustering of the same ANI matrix. Secondly, by constructing a maximum likelihood phylogeny inferred with GToTree from the Actinobacteria single-copy marker-gene set (Lee, 2019). This tree was midpoint-rooted and species were grouped by slicing at 70% of the maximum root-to-tip depth, taking the resulting monophyletic clades as higher-order complexes (Supplementary Figure 2).

### Pangenome analysis

Pangenomes were computed per species following de novo species designation, using quality-filtered genomes retrieved from NCBI Datasets and our clinical isolate genomes. Species were included in pangenome analysis if they had at least 20 genomes and contained >1 isolate from our cohort. For each species, PPanGGOLiN was used to infer gene families, partitioning the species gene repertoire into the core (present in ≥ 99% of genomes), shell (intermediate-frequency) and cloud (rare or strain-specific) (Gautreau et al., 2020). To evaluate pangenome openness, gene-accumulation curves were averaged over 50 random genome-input orders, rarefied to a common depth of 20 genomes and fitted to Heaps’ law (Tettelin et al., 2008, 2005). A 95% confidence interval on the accumulation exponent (α) was obtained by bootstrapping genomes (300 resamples). Species were grouped into tiers by 95% confidence interval overlap. Species whose intervals did not overlap were regarded as distinguishable, and the number of distinguishable species pairs was quantified. Genome fluidity, which describes the mean fraction of unshared gene families between genome pairs, was used to summarise accessory-genome divergence. In the latter case, 95% confidence intervals were calculated from a delete-one jackknife, with species again grouped into tiers by confidence-interval overlap using the same procedure (Kislyuk et al., 2011). The concordance between openness and fluidity was assessed with Spearman’s rank correlation.

### Genome annotation, mobilome and clinical-isolate comparisons

Bakta was used for protein prediction and genome annotation, with unannotated proteins rescued via Baktfold (Bouras et al., n.d.; Schwengers et al., 2021). Predicted protein sequences produced by Bakta were used for downstream functional annotation via multiple methods. EggNOG-mapper was used for gene level functional annotation against the eggNOG database (Cantalapiedra et al., 2021). KEGG Orthology (KO) assignments were made by running KofamScan against the KOfam profile HMMs with default per-KO adaptive thresholds (Aramaki et al., 2020; Kanehisa et al., 2023). Carbohydrate active enzymes were annotated using dbCAN, retaining calls with a consensus across at least two of its three methods (HMMER, dbCAN-sub, DIAMOND) (Zheng et al., 2023). DbCAN-sub eCAMI subfamily identifiers were collapsed to their parent CAZy family using subfamily suffixes before counting. Secondary-metabolite biosynthetic gene clusters were annotated by antiSMASH (Blin et al., 2023). KEGG module completeness was assessed per genome; completeness was taken as the fraction of constituent KO steps present, and a module was called complete where ≥80% of its steps were present. Feature prevalence was calculated as the fraction of isolates in which the feature was encoded (KEGG module complete, CAZyme family present, or BGC class detected). Species-specific enrichment was tested by one-versus-rest permutation testing (20,000 permutations, Benjamini-Hochberg FDR correction q<0.05) and only applied to species with ≥ 4 clinical isolates. Significant features were visualised where they fell into the cohort-variable set or if they were cohort-rare but substantially complete in the species that carried them compared to rest of cohort (in-group mean completeness ≥0.5, outgroup <0.3).

Two methods of antimicrobial resistance gene annotation were applied; AMRFinderPlus and CARD/RGI (Alcock et al., 2023; Feldgarden et al., 2021). Virulence genes were annotated by matching against VFDB with DIAMOND (Buchfink et al., 2021; Liu et al., 2022). Annotation hits were retained where they met tool-specific thresholds: for VFDB, ≥70% identity, ≥70% query coverage and E-value ≤1×10^-10^; for AMRFinder ≥70% identity, ≥70% query coverage and exclusion of partial hits; and for RGI inclusion of only perfect or strict matches with ≥70% identity, and ≥70% reference length. A final consensus annotation for virulence and AMR was then made per protein with VFDB taking precedence over CARD/RGI and AMRFinderPlus filling in the residual. Virulence-factor status required a coding sequence’s own VFDB alignment, not gene-symbol identity with a VFDB entry, since VFDB also catalogues conserved housekeeping genes. Symbol matching would have returned 1,095 loci against the 834 supported by alignment. For virulence, annotation reliability was assessed per clade using a positive control panel of essential, genus-conserved genes (GroEL and EF-Tu) to account for poor representation of commensal species within the VFDB.

Metabolic pathways, CAZymes, BGC classes, virulence factors and AMR determinants were aggregated for comparison of genome functional capacity at the species and strain level. Inter-species comparisons were made across the clinical isolate set only. To describe the gene repertoire of the cohort sinus-isolates, annotations were partitioned into a cohort-shared set (carried by ≥95% of isolates), a cohort-variable set (5 to 95%), and a cohort-rare set (<5% carriage).

Mobile genetic elements were identified using the European Bioinformatics Institute (EBI) Mobilome Annotation Pipeline (v4.2.3; geNomad, ICEfinder2, IntegronFinder, ISEScan) to place the Bakta gene annotations in their mobile-element context. Each coding sequence was assigned to a mobile-element compartment (chromosomal, insertion sequence, plasmid, prophage) based on the overlap between its locus and the mobilome consensus annotation. Mobilome architecture was reported as the median number of coding sequences per genome in each compartment. Virulence and AMR determinants were localised by compartment, and non-chromosomal assignments were reported. All per-genome processes were parallelised using GNU parallel (Tange, 2018). Using a two-sided exact binomial test, compartment enrichment was assessed per determinant class by comparing the fraction of its loci with the share of all coding sequences in the same genomes; 95% confidence intervals on the fold difference were taken as the Clopper-Pearson interval of the locus rate divided by the background rate. For this analysis the horizontally acquired compartment pools plasmid, prophage, integron and integrative-conjugative-element assignments and excludes insertion sequences.

## 3. Results

### Genome collection and clinical cohort

In this study we have analysed the genomes of 87 *Corynebacterium* clinical isolates, collected from the sinonasal cavity of 71 patients. This Australian-based cohort comprised 20 non-CRS control; 25 CRS without nasal polyp; and 26 CRS with nasal polyp patients, contributing 26, 31 and 30 isolate genomes respectively (Table 1). Clinical isolates were first speciated by matrix-assisted laser desorption/ionisation time-of-flight mass spectrometry (MALDI-TOF MS), before final species assignment by whole genome sequencing.

**Table 1.**
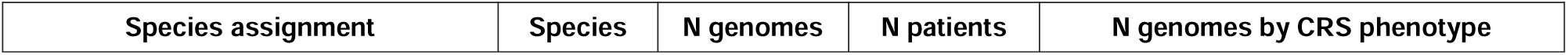
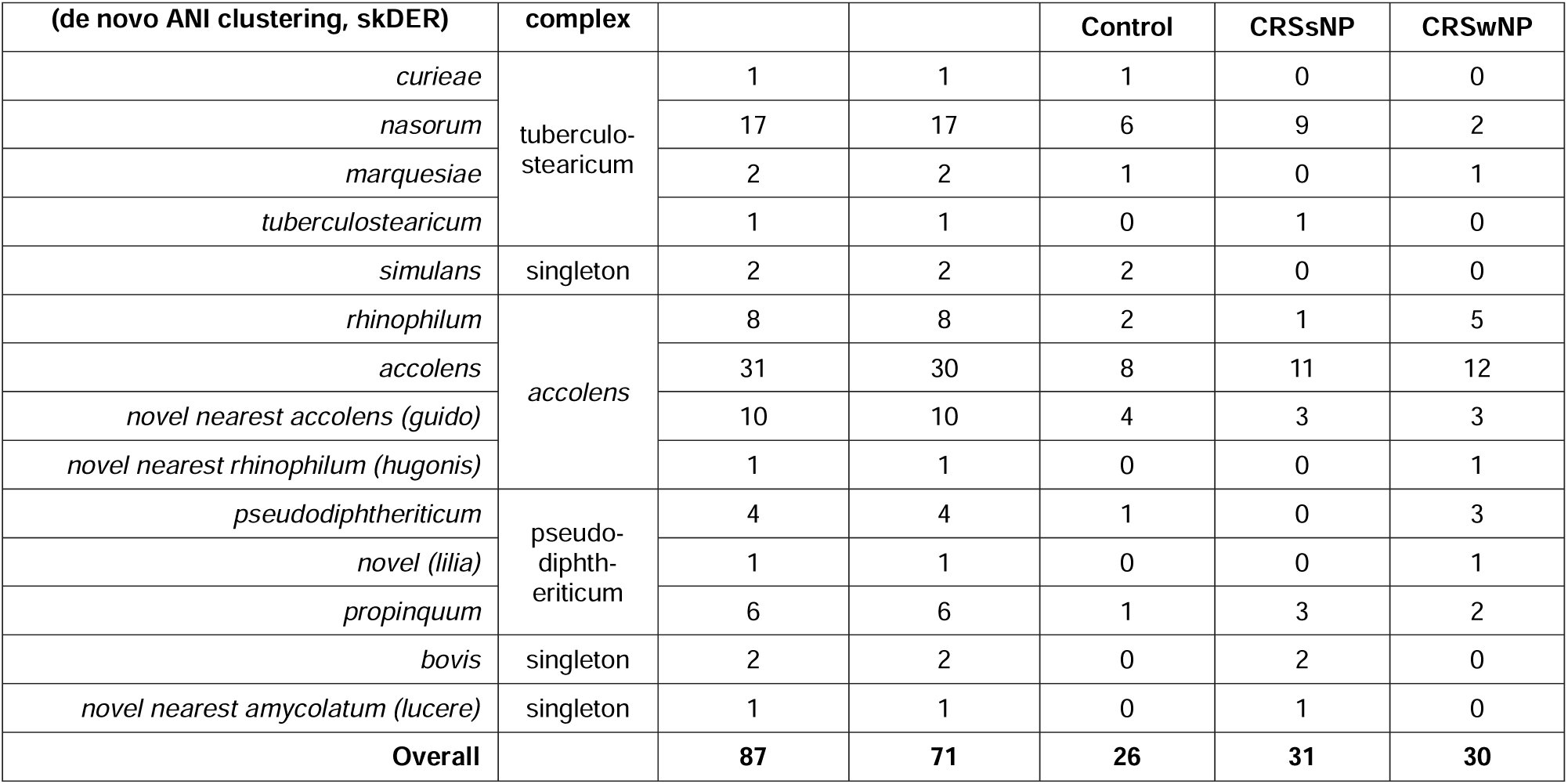
Number of clinical isolates in the final cohort by species and clinical phenotype.

Final genome-based skDER species assignment agreed with MALDI-TOF MS results at the species-complex level (Table 1; Figure 1A,D). At the species level, whole genome sequencing with skDER species assignment reassigned 41 of 87 isolates relative to their MALDI-TOF MS identification (Figure 1A). Species assignments by skDER were broadly concordant with GTDB-Tk, as expected given the shared ANI-based approach (Figure 1B) (Chaumeil et al., 2022; Parks et al., 2022; Salamzade et al., 2025). Residual differences reflect the curated LPSN- and literature-based reference set used to anchor species names in this study (Figure 1B) (Parte et al., 2020; E. B. Popowitch et al., 2024). Four candidate novel species were identified and assigned the following placeholder names; *Corynebacterium guido* (novel taxon, nearest *C. accolens*)*, Corynebacterium hugonis* (novel taxon, nearest *C. rhinophilum*)*, Corynebacterium lucere* (novel taxon, nearest *C. amycolatum*)*, and Corynebacterium lilia* (novel taxon, no species within 90% ANI). The genome size of our corynebacteria ranged from 2.3 Mbp to 2.7 Mbp (average 2.5 Mbp), a minimum completeness of 99.9% (median 100%) and maximum contamination of 0.66% (median 0.06%) (Figure 1E). Unlike other *Corynebacterium* species in the cohort, *C. bovis* had a median GC content of 73%. Excluding *C. bovis*, median GC content for genomes in the dataset was 59% with a minimum of 55% and max of 60% (Figure 1E).

**Figure 1.**
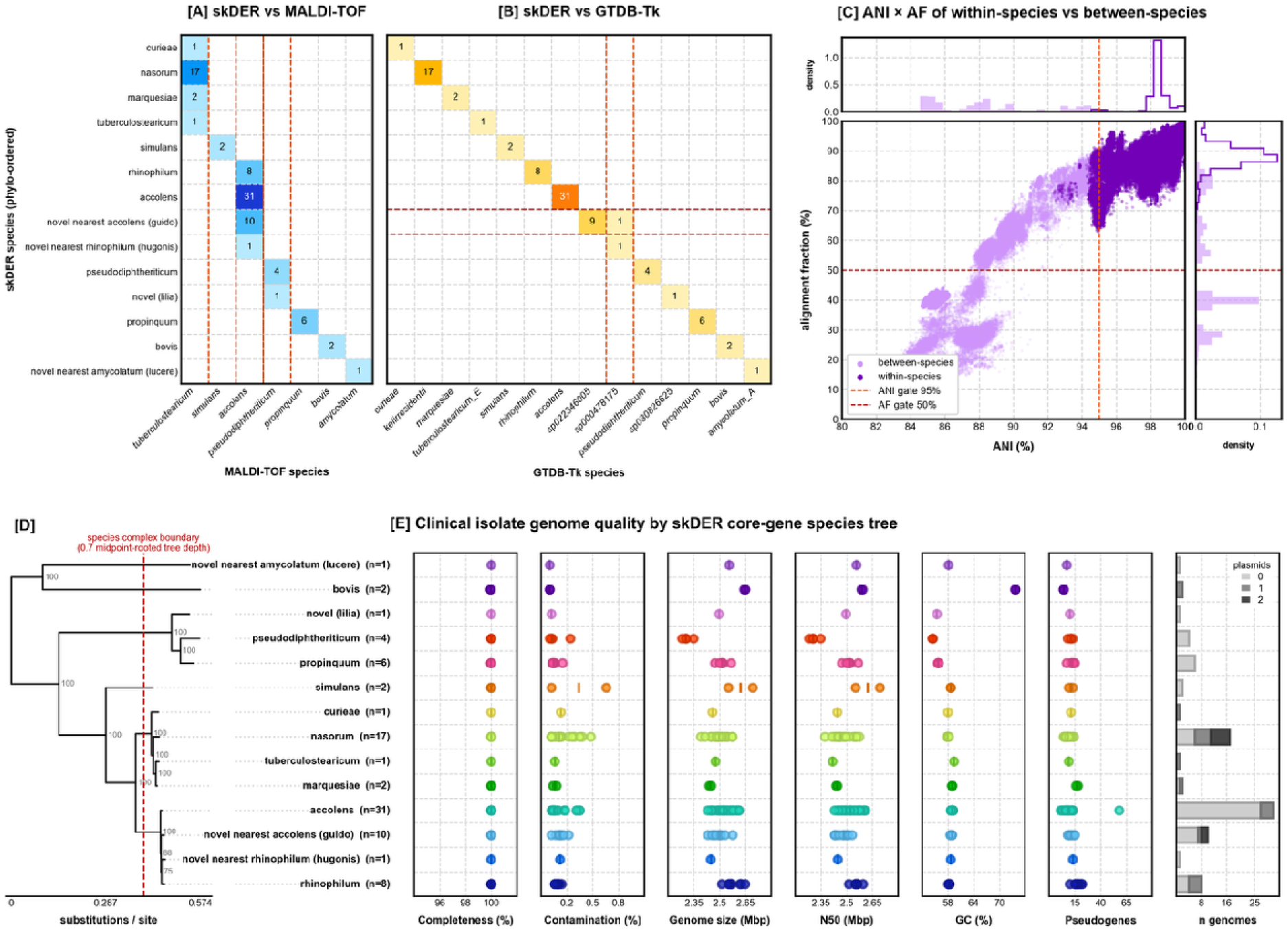
Representation of final species assignments, phylogeny and genome quality in the final cohort. [A] species classification by skDER vs MALDI-TOF MS; [B] species classification by skDER vs GTDB-Tk; [C] whole-Corynebacterium-genus all versus all average nucleotide identity (ANI) comparison and alignment fraction (AF) of 4661 quality filtered Corynebacterium genomes accessed via NCBI Datasets and respeciated by skDER; [D] maximum-likelihood marker-gene tree of the 87 clinical isolate genomes (GToTree, Actinobacteria single-copy gene set), midpoint-rooted and collapsed to one leaf per skDER species; node labels are bootstrap support and the dashed line is the species-complex boundary at 0.7 of the midpoint-rooted tree depth; [E] completeness, contamination, genome size, N50, GC content, number of pseudogenes and number of plasmids per genome (stacked bars) in the clinical isolate genomes.

### The *Corynebacterium* landscape and metadata review

All *Corynebacterium* genomes accessible via NCBI Datasets were downloaded together with their metadata yielding 5,055 assemblies, of which 4,661 passed quality filtering for speciation. The deposition of Corynebacterium genomes has roughly been increasing year on year with 808 genomes deposited in 2025 and a further 812 in 2026 to the access date (Figure 2C). Of the existing *Corynebacterium* assemblies, only 731 were complete genomes or chromosomes (Figure 2A). The majority of these genomes have been isolated from human hosts (71.1% of known hosts, Figure 2B), in particular, the human respiratory tract (41.7% of known body sites, Figure 2D). The next most common sites of *Corynebacterium* isolation were wound/abscess, nasal/sinus, blood and skin (8.7%, 8.4%, 6.4%, 6.1% respectively, Figure 2D). Published *Corynebacterium* genomes predominantly originated from the United States of America and China (863, 853 genomes respectively), with only 64 published Australian *Corynebacterium* genomes (Figure 2E).

**Figure 2.**
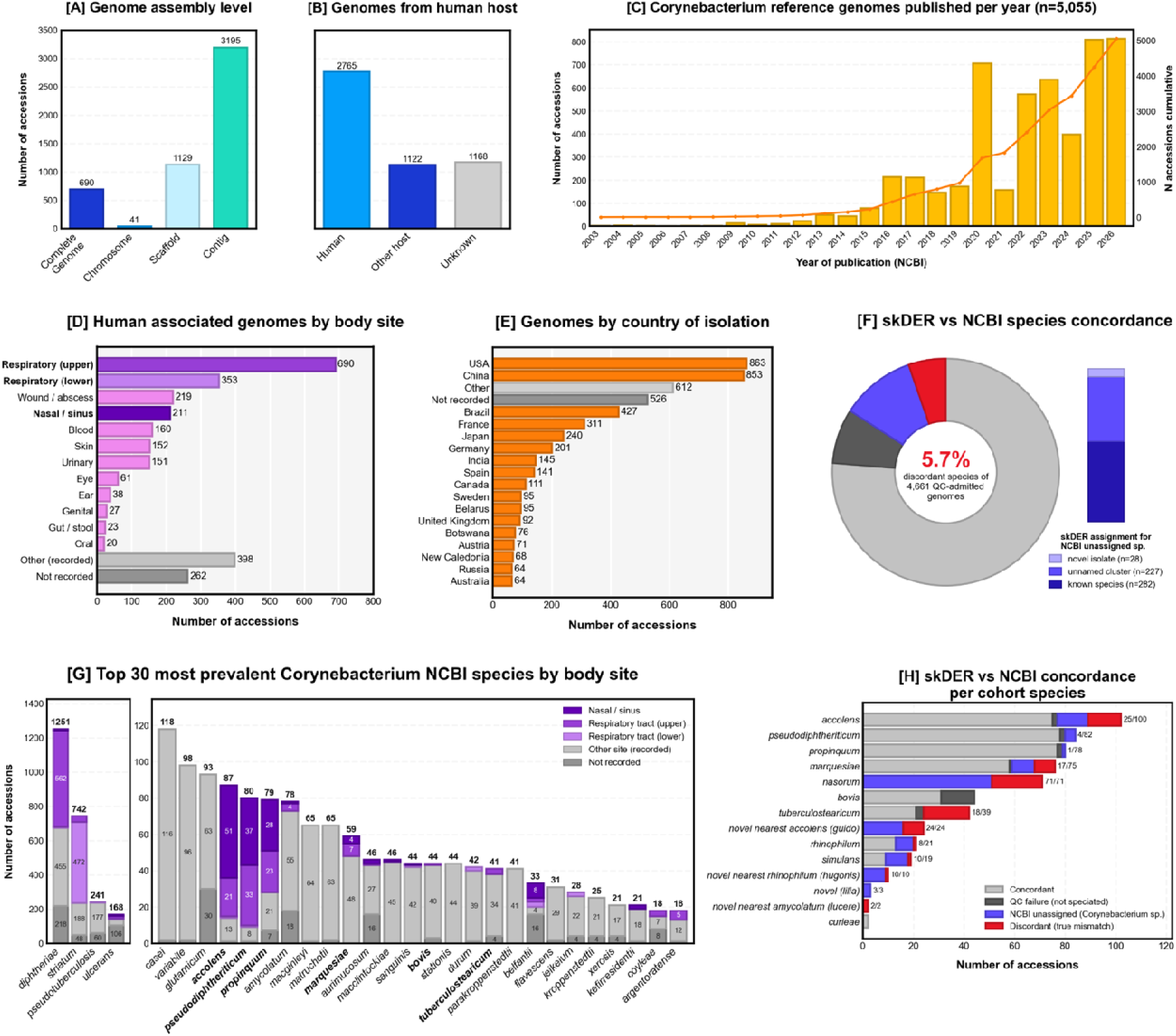
Metadata review of Corynebacterium genomes accessed via NCBI Datasets (accessed 8 August 2026). [A] assembly level of published Corynebacterium genomes; [B] number of Corynebacterium genomes with human-host isolation source; [C] annual rate of Corynebacterium deposition to NCBI Datasets; [D] human-isolated Corynebacterium genomes enumerated by body site; [E] genomes enumerated by country of isolation (top 17 countries); [F] donut chart showing the proportions of skDER species assignments concordant with NCBI taxonomy, NCBI genomes failing QC, NCBI accessions with no assigned species, discordant species, and stacked bar representing outcome of respeciation for NCBI accessions with no assigned species; [G] most prevalent Corynebacterium species coloured by body site of isolation; [H] skDER and NCBI taxonomy concordance within the taxa included in the current clinical isolate cohort.

The best represented *Corynebacterium* species were *C. diphtheriae,* the causative agent for diphtheria, *C. striatum*, and the zoonotic pathogen, *C. pseudotuberculosis* (accounting for 24.7%, 14.7%, 4.8% of *Corynebacterium* genomes respectively; Figure 2G). These species were rarely represented by isolates from nasal/sinus sites (Figure 2G). The most well sequenced sinus-associated *Corynebacterium* species were *C. accolens* (87 genomes, 59% nasal/sinus), *C. pseudodiphtheriticum* (80 genomes, 46% nasal/sinus), and *C. propinquum* (79 genomes, 35% nasal/sinus; Figure 2G).

Among the 4,661 genomes that passed quality filtering, de novo species assignment by skDER was broadly concordant with the species names recorded in NCBI Datasets (82.8% concordant; Figure 2F). Of the residual, 5.7% (n=266) were assigned to a different species, whilst 11.5% were genomes with no NCBI species-level label. Amongst the latter 537 accessions lacking an NCBI species designation, skDER assigned 282 genomes to a known species; 227 fell in clusters with no type strain; and 28 genomes found a species level match with the novel species defined in this cohort. In total, 39 public genomes fell within the four candidate novel species boundaries, the aforementioned 28 unnamed genomes, as well as 11 genomes deposited to NCBI labelled as a different species (*C. guido*, n=24; *C. hugonis*, n=10; *C. lilia*, n=3; *C. lucere*, n=2) (Figure 2F). NCBI-skDER concordance for the species represented in this sinus-isolate cohort is shown in Figure 2H. Species re-assignment was frequent for *C. amycolatum* (72 reassigned / 145 NCBI assemblies), *C. accolens* (25/100), *C. tuberculostearicum* (18/39), *C. marquesiae* (17/75), and *C. simulans* (10/19), but rare for *C. propinquum* (1/78) and *C. pseudodiphtheriticum* (4/82) (Figure 2H). All 71 public genomes of the *C. nasorum* lineage were reassigned, 51 from no species designation, and 18 from *C. kefirresidentii*, and one each from *C. tuberculostearicum* and *C. aurimucosum* (Figure 2H).

### Pangenomes

Species pangenomes were calculated from all available genomes (NCBI plus study isolates) for each of the species in this cohort represented by ≥20 genomes (n=10; Figure 3A). Core gene family counts were similar across the tested species, ranging from 1,597 (*C. marquesiae*) to 1,878 (*C. guido*). Core content as a proportion of the pangenome varied from 28% (*C. marquesiae*) to 57% (*C. guido*) (Figure 3A). Pangenomes were open for all species, with no plateau reached in the accumulation curves (Heaps’ α>0) (Figure 3B,C). However, the extent of openness varied. When rarefied to a common depth of 20 genomes for comparability, openness ranged from α_20_=0.095 (*C. guido*) to α_20_=0.195 (*C. marquesiae*) (Figure 3D). The three least open species were C*. guido, C. bovis* and *C. rhinophilum*; (α_20_ 0.095-0.102), their confidence intervals overlapped with one-another but no other species. Genome fluidity delineated the accolens complex more clearly; the confidence intervals in these species were non-overlapping with that of the top tier (Figure 3E). Pangenome openness and fluidity were concordant (Spearman ρ=0.88, p=0.001; Figure 3F), with *C. rhinophilum* and C*. guido* at the lower end, and *C. pseudodiphtheriticum*, *C. simulans* and *C. marquesiae* clustered at the higher end.

**Figure 3.**
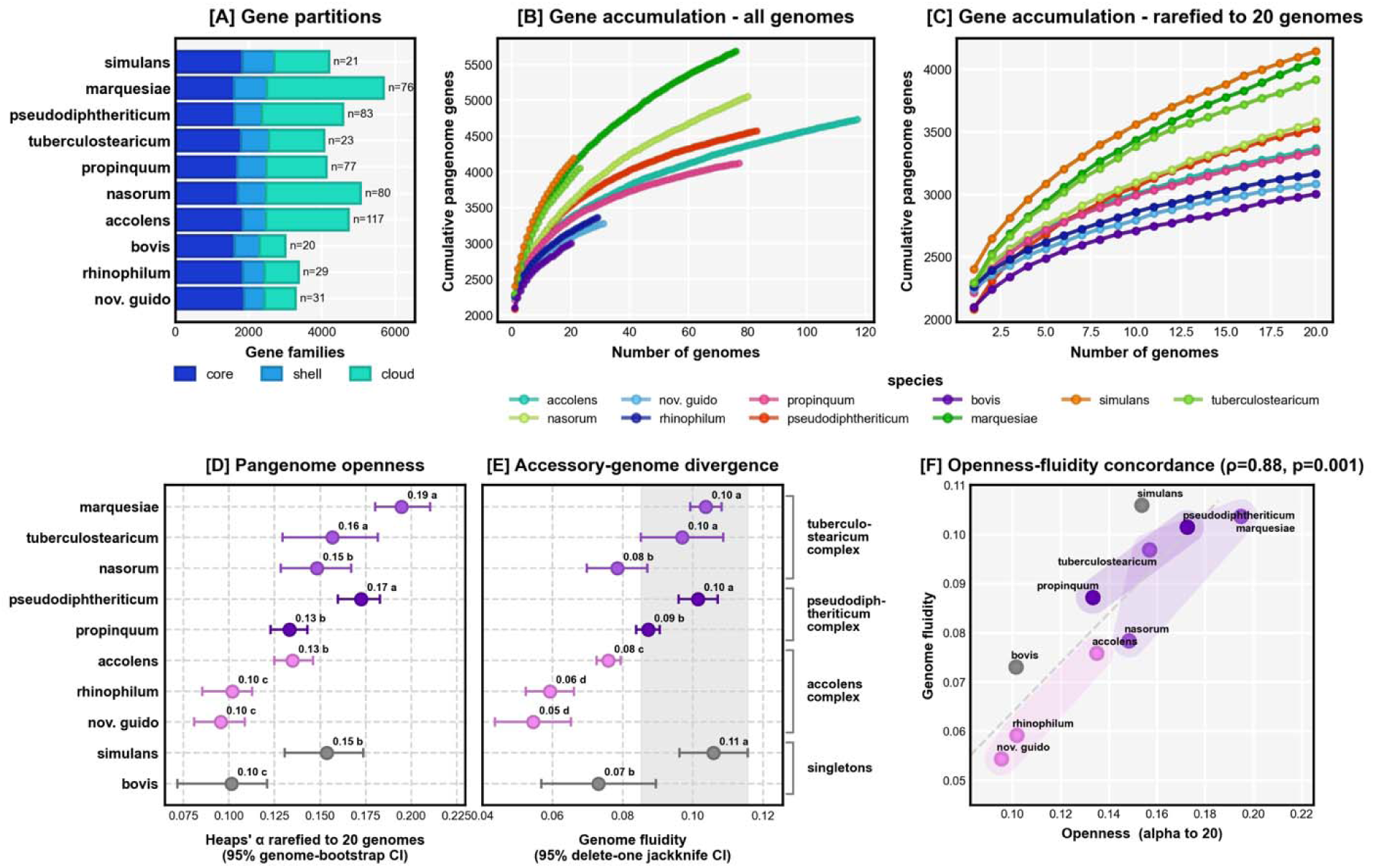
Species pangenome repertoires and dynamics derived using PPanGGOLiN [A] size of core, shell and cloud gene repertoire per species (n genomes per pangenome); [B] species pangenomes calculated using all available genomes (NCBI and clinical isolates); [C] species pangenomes in well-sampled species (min strains n=20) rarefied to 20 genomes; [D] median Heaps’ accumulation exponent (α) and 95% confidence intervals (bootstrapping 300 resamples) plotted per species, rows are grouped by species complex and ordered by decreasing median α within each complex (singletons last), the letter annotating each point represents its tier by CI overlap; [E] median genome fluidity and 95% confidence intervals (95% delete-one jackknife) in the row order of [D], the grey band spans the CIs of the top tier (tier A); [F] pangenome openness (Heaps’ α rarefied to 20 genomes) plotted against genome fluidity, dashed line shows linear fit [B,C] Point estimates coloured by species [D-F] Point estimates coloured by species complex (singletons grey).

### Function and metabolism

Encoded functions were broadly conserved across sinus *Corynebacterium* isolates, with a subset of variable features separating along both species-complex and species boundaries (Figure 4). Complete KEGG modules for amino acid and carbohydrate metabolism were largely shared (80% and 61% respectively), whereas cofactors were more divergent, with only 47% of complete modules shared by ≥95% of isolates (Figure 4B). Lipid and terpenoid metabolism each contributed only three complete modules and varied with phylogenetic lineage: C10-C20 isoprenoid biosynthesis (M00364/M00365) was complete in every isolate except *pseudodiphtheriticum*-complex genomes (0/11) and *C. bovis* (0/2). The fatty acid biosynthesis module M00083 showed the reverse pattern, complete in all *pseudodiphtheriticum*-complex genomes (11/11) and singletons *C. lucere* (1/1) and *C. simulans* (1/2) but absent from every *accolens*- and *tuberculostearicum*-complex genome. This is consistent with the lipophilic growth requirements characteristic of these lineages, implying dependence on host-derived lipid (Figure 4B,E). At the gene level, the lipase *lipS1* and esterase *estA1* were near exact complements: *lipS1* was carried by all *pseudodiphtheriticum*-complex (11/11) and nearly all *tuberculostearicum*-complex genomes (20/21) but was absent from *accolens*-complex genomes (0/50); meanwhile, genomes within the *accolens*-complex universally encoded the *estA1* (50/50 genomes), a gene absent from *pseudodiphtheriticum*-complex genomes (0/11) and, amongst the *tuberculostearicum*-complex, only encoded by select *C. nasorum* strains (13/17) (Supplementary Figure 3).

**Figure 4.**
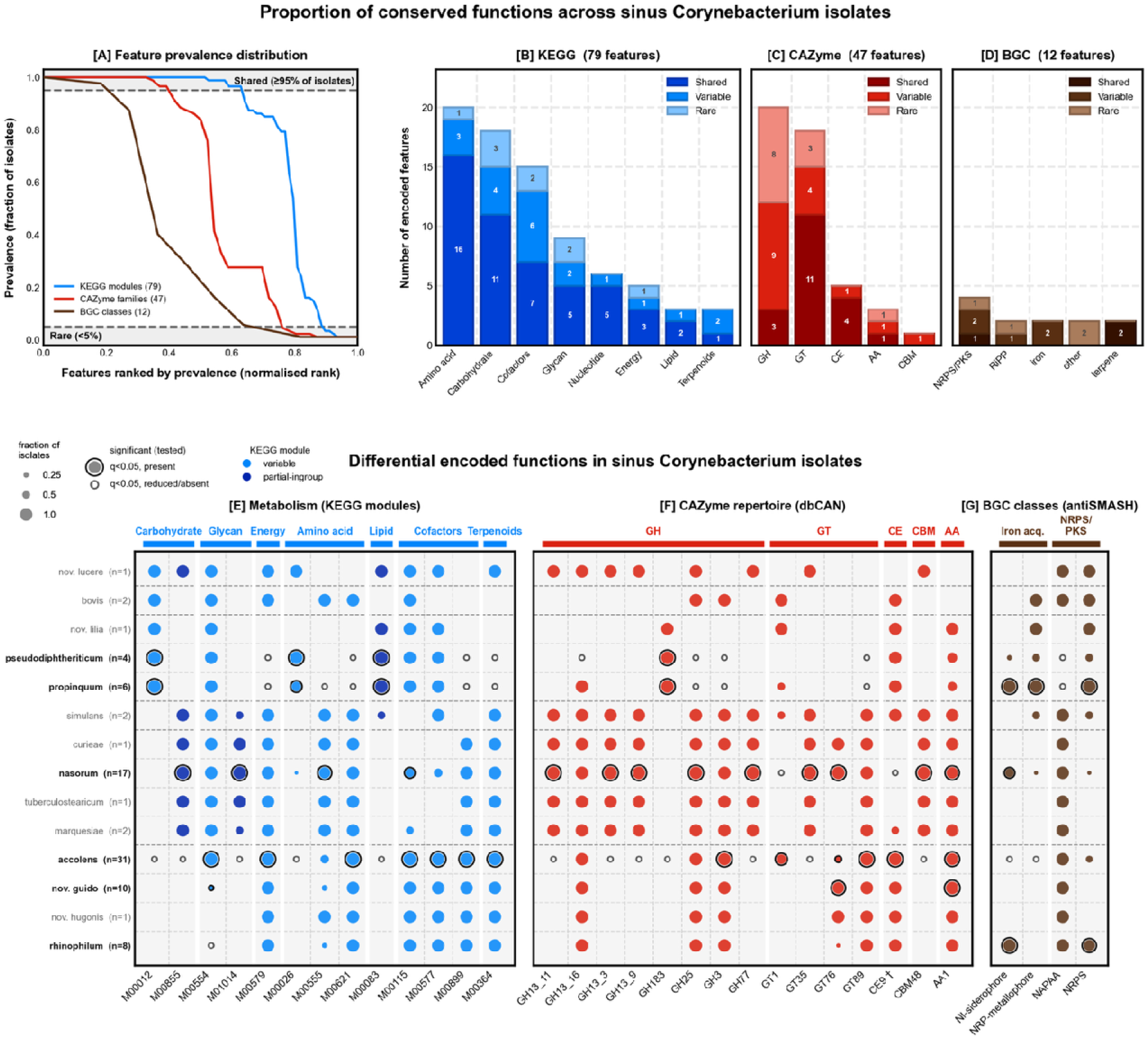
Functional profiles of sinus Corynebacterium isolates [A] prevalence of complete KEGG modules, CAZyme families and BGC classes in clinical isolates; [B-D] number of shared (feature present in ≥95% clinical isolates), variable, and rare (<5%) complete KEGG modules, CAZymes and biosynthetic gene clusters (BGC); [E-G] fraction of isolates encoding differential completed KEGG modules, CAZymes and BGCs displaying significantly differential features from the variable compartment, except dark blue KEGG modules which were rare across the cohort but common to a species complex. Circle size is the fraction of isolates in each species, and species with fewer than four isolates are shown but were not tested.

Amongst CAZymes, carbohydrate esterase (CE) and glycosyltransferase (GT) repertoires were the most conserved (80% and 61% shared respectively). By contrast, glycoside hydrolases (GH) were highly variable, with only 15% shared features and 40% rare features (Figure 4C). Carbohydrate acquisition split these sinus corynebacteria lineages, at a complex level for glycogen, and a species level for sialic acid. The glycogen utilisation CAZymes (GH13_3, GH77 and GT35), as well as the glycogen-binding module (CBM48), were carried by every member of the *tuberculostearicum*-complex (21/21) and no accolens-complex genome (0/50) (Figure 4E, F). Similarly, glycan degradation (M00855) and UDP-ManNAcA biosynthesis (M01014) were largely restricted to the *tuberculostearicum*-complex (21/21 and 20/21 genomes). The only other carriers were the singletons *C. simulans* (2/2) and *C. lucere* (1/1) (Figure 4E). As an alternate source of carbohydrates, the *pseudodiphtheriticum*-complex instead carried the sialic acid catabolic operon *nanAEK* across all genomes (11/11), as did a subset of *C. accolens* isolates (8/31) (Supplementary Figure 3). Within the *accolens*-complex there was a loss of UDP-Gal biosynthesis (M00554) in *C. rhinophilum* (0/8 genomes), *C. hugonis* (0/1), and *C. guido* (1/10), despite its retention in all *C. accolens* isolates (31/31) (Figure 4E). The *pseudodiphtheriticum*-complex was broadly CAZyme depleted; however, *pseudodiphtheriticum*- and *accolens*-complex genomes encoded the host-glycan-active esterase CE9 (11/11 and 50/50 respectively) which was largely absent from the *tuberculostearicum*-complex (1/21) (Figure 4E, F).

Anaerobic nitrate respiration further distinguished species complexes, with presence of the nitrate-reductase operon (*narGHIJ*) and nitric oxide reductase (norB) across all *accolens*- and *pseudodiphtheriticum*-complex genomes (50/50 and 11/11) but largely absent throughout the *tuberculostearicum*-complex (narGHIJ 1/21 and norB 0/21), indicating loss of anaerobic nitrate respiration in that lineage (Supplementary Figure 3). At the species level, genes encoding the flavohaemoglobin *hmp* and its regulator *nsrR* were restricted to *C. pseudodiphtheriticum* (4/4), *C. bovis* (2/2), and *C. lucere* (1/1) (Supplementary Figure 3). Similarly, a complete urease operon (*ureABCDG*) was largely confined to *C. bovis* (2/2) and *pseudodiphtheriticum*-complex (10/11) genomes, providing a route to nitrogen salvage and local alkalinisation (Supplementary Figure 3). The glyoxylate cycle (M00012) was confined to the *pseudodiphtheriticum*-complex (11/11) and singletons *C. bovis* (2/2) and *C. lucere* (1/1) and absent from every *accolens*-(0/50) and *tuberculostearicum*-complex (0/21) genome (Figure 4E). Conversely, accolens- and tuberculostearicum-complex genomes shared the phosphotransacetylase-acetate kinase pathway (M00579; 71/71 genomes), which the *pseudodiphtheriticum*-complex lacked (0/11 genomes), indicating divergent strategies for acetate handling (Figure 4E).

Cofactor biosynthesis likewise varied by lineage: de novo biotin biosynthesis (M00577) was complete in all *accolens*- and *pseudodiphtheriticum*-complex genomes, and least complete in the *tuberculostearicum*-complex (10/21, all of them *C. nasorum*) and absent from C. bovis (0/2) (Figure 4E). Genomes of the *accolens-* and *tuberculostearicum*-complexes encoded a capacity for thiamine salvage (M00899;71/71), absent from the *pseudodiphtheriticum-*complex (0/11) and five singleton genomes (Figure 4E).

BGCs were the most sparsely represented feature set with 12 classes detected, of which 25% were shared, 42% variable and 33% rare (Figure 4D). Non-alpha poly-amino acid clusters (NAPAA) were encoded by most isolates (76/87) but were universally absent from the *pseudodiphtheriticum*-complex (0/11) (Figure 4G). This complex was instead enriched in non-ribosomal peptide synthetase (NRPS) and NRP-metallophore clusters, particularly in *C. propinquum* (6/6 isolates for each). Iron acquisition varied within the *accolens*-complex: *C. rhinophilum* encoded NRPS-independent siderophores (8/8) which were absent from every other accolens-complex genome (0/42) (Figure 4G).

*C. rhinophilum* also encoded NRPS cluster (7/8), which was carried by only 13/31 *C. accolens* isolates and absent from *C. guido* and *C. hugonis* (Figure 4G).

### Virulence factors and Antimicrobial Resistance

In the sinus *Corynebacterium* isolates (n=87), 42 virulence factors and 7 acquired antimicrobial resistance genes were detected (Figure 5A, B). Under the alignment-based criterion GroEL (groEL2) was detected in 87 of 87 genomes and EF-Tu (tufA) in 72 of 87 (83%), confirming that VFDB detection held across the cohort. The prevalence of virulence and AMR was low, contributing to 834 (VF) and 62 (AMR) loci out of a total of 202,706 coding sequences across the cohort. Every isolate carried at least one virulence factor, whilst 36 of 87 carried at least one acquired resistance gene. Encoded capacity for virulence and antimicrobial resistance varied by species (Figure 5C). AMR gene prevalence was least in the *accolens*-complex (17% mean prevalence across drug classes), followed by the *tuberculostearicum*-complex (26%), and greatest in the *pseudodiphtheriticum*-complex (31%). These most commonly encoded resistance to streptogramin, lincosamides and macrolides (Figure 5E). Among well sampled species (n≥4), virulence factor density (median VF genes per 1k CDS) was lowest in *C. guido* (2.6), *C. nasorum* (3.0), and *C. rhinophilum* (3.4), and greatest in *C. propinquum* (5.5) and *C. pseudodiphtheriticum* (5.8) (Figure 5C, D).

**Figure 5.**
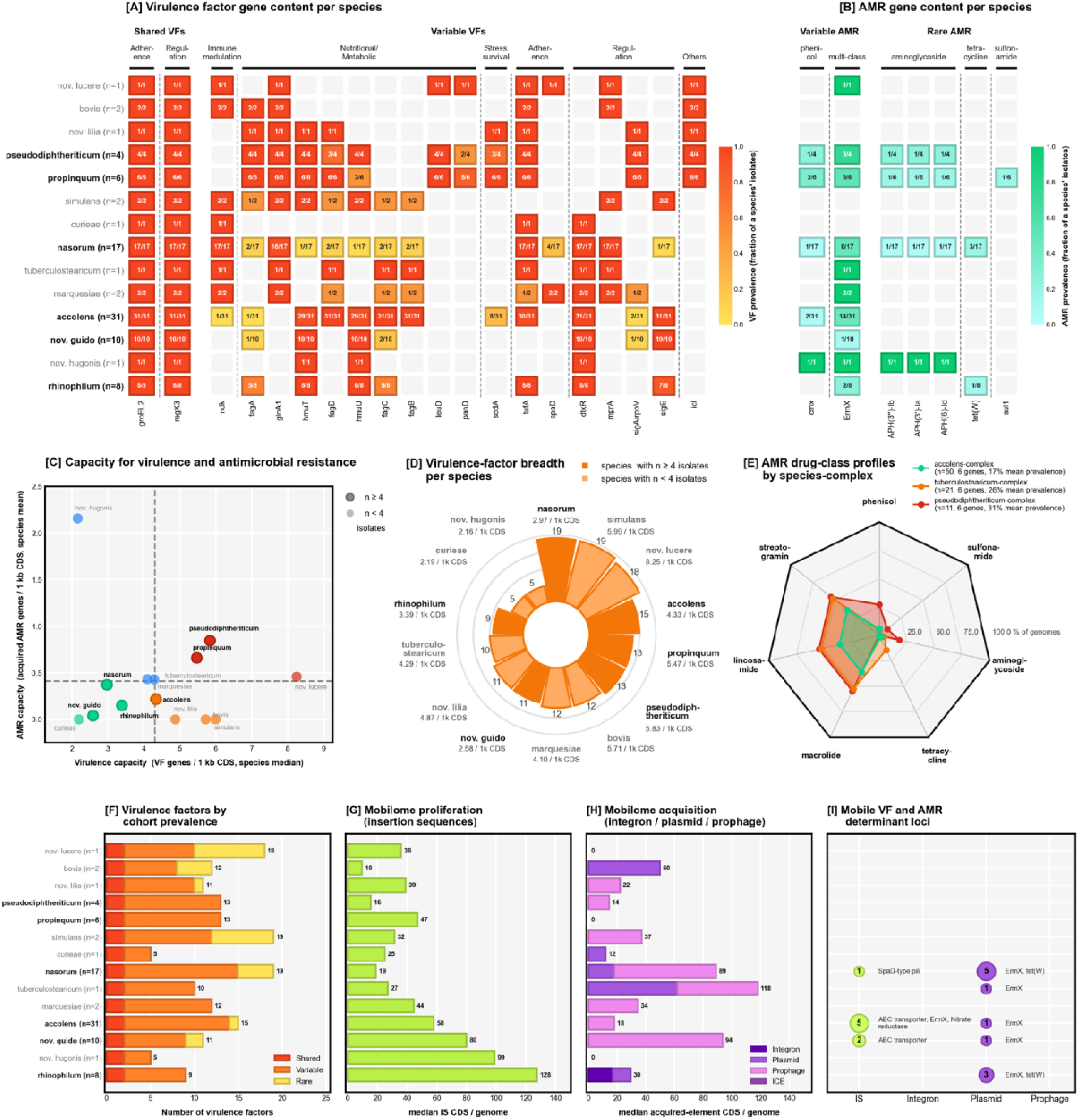
Sinus Corynebacterium virulence and antimicrobial resistance repertoire, and context within mobilome architecture, species y axes ordered by phylogeny [A] prevalence of shared, variable and rare virulence factors by species; [B] prevalence of variable and rare AMR genes by species (shared AMR n=0); [C] species median virulence factor (VF) density plotted against species mean AMR gene density; [D] breadths and median density of virulence factors by species; [E] prevalence of AMR genes grouped by drug class, per species-complex; [F] number of virulence factors in shared (common to ≥95% of genomes), variable and rare (<5% of genomes) compartments by species; [G] median number of insertion-sequence (IS) coding sequences per genome plotted by species; [H] median number of integron, plasmid and prophage coding sequences per genome plotted by species; [I] number of AMR and VF determinant loci residing within mobile-genetic elements by species; each bubble represents the total number of loci in that compartment and lists the VF and AMR genes.

Virulence factor breadth and VF density diverged in the case of *C. nasorum* which demonstrated the greatest number of distinct VFs per species (n=19 VFs across 17 isolates); of the well-sampled species, *C. accolens* had the next greatest VF breadth carrying 15 VFs across 31 isolates (Figure 5D). Two virulence factors were shared across sinus *Corynebacterium* isolates (≥95% of isolates), 18 were variable and 22 were rare, with 17 of these rare factors confined to a single species (Figure 5A, F). This variation was not evenly distributed but driven by specific virulence categories (Figure 5A). The two shared virulence factors were housekeeping genes carried by every isolate: GroEL (groEL2) and the RegX3 response regulator. Elongation factor Tu (tufA, 83% of isolates) and the iron-dependent regulator DtxR (82%) fell just below the shared threshold, followed by haem-uptake HmuTU (75%, 65/87 genomes) (Figure 5A). Targeted screening for the canonical *C. diphtheriae* toxin gene (tox) found universal absence in this cohort. Variable virulence factors were largely partitioned by species-complexes (Figure 5A). The *tuberculostearicum*-complex, captured the greatest breadth of regulatory virulence factors (Figure 5A; Supplementary Figure 3). The *pseudodiphtheriticum*-complex encoded pantothenate synthesis (*PanD*) and isocitrate lyase (*icl*) genes, absent from other species-complexes, consistent with the greater completeness of the glyoxylate cycle in this complex (M00012; Figure 4E, Figure 5A).

Iron acquisition machinery was encoded near-universally across sinus corynebacteria. Every isolate encoding the ferric-hydroxamate uptake ATPase gene *fhuC* (Supplementary Figure 3). However, several uptake systems diverged at the species and species-complex level. The haem uptake *hmuTU* was carried by the majority of *accolens*-(49/50) and *pseudodiphtheriticum*-complexes (7/11, absent from *C. propinquum*), but only present in one *C. nasorum* isolate amongst the *tuberculostearicum*-complex (Figure 5A). Select features of the Fag hydroxamate uptake system (*fagABCD*) were encoded across the *pseudodiphtheriticum-*complex (*fagA* 11/11; *fagD* 10/11) but varied across *C. accolens* isolates (*fagBCD* 31/31; *fagA* 1/31), *C. rhinophilum* isolates (*fagAC* 5/8), and *C. guido* isolates (*fagC* 2/10; *fagA* 1/10), and was largely absent in the *tuberculostearicum*-complex (*C. nasorum fagABCD* 2/17, *C. marquesiae fagBCD* (1/2), *C. tuberculostearicum fagBCD* 1/1) (Figure 5A). The iron-dependent regulator dtxR was also split across lineages, present in every *accolens*- and *tuberculostearicum*-complex genome (50/50, 21/21) and absent from the *pseudodiphtheriticum*-complex (0/11) and all singletons.

Mobile-element classification assigned 4.74% of coding sequences (9,599 of 202,706 CDS) across 87 genomes to a mobile compartment. The mobile compartment was further split into insertion sequences (IS, 2.42% of CDS), indicative of intragenomic proliferation, and horizontally acquired elements (integrons 0.09%; plasmids, 0.52%; and prophages, 1.7% of CDS). The contents of the mobilome varied at a species and strain level (Figure 5G, H, Supplementary Figure 4). The cohort-wide median number of insertion sequences was 37 and, in well sampled species (n≥4), ranged from 15.5 in *C. pseudodiphtheriticum* to 127.5 in *C. rhinophilum* (Figure 5G). The median number of horizontally acquired genetic elements was also highly variable (range 0 to 118; Figure 5H). Prophages were the most commonly observed acquired element, followed by plasmids, with integrons the smallest compartment throughout (Figure 5G, Supplementary Figure 4).

AMR determinants were significantly enriched on plasmids. Eleven of 62 resistance loci (17.7%) resided in this compartment, which comprised only 0.52% of all coding sequences, a 34-fold enrichment (95% CI 17.6 to 56.4, exact binomial p=3.2×10^-14^). This signal was driven entirely by plasmids, which carried all 11 loci, despite only representing 0.52% of all CDS, culminating in a 34-fold enrichment of AMR genes on plasmids (95% CI 17.6 to 56.4, p=3.2×10^-14^; Figure 5H, I). Prophages and integrons did not encode AMR; and whilst two AMR genes were detected on insertion sequences, plasmids remained the only mobile compartment to demonstrate this significant relationship (3.2% of AMR loci on IS against 2.4% of CDS; exact binomial p=0.66). The remaining 49 resistance loci were chromosomal. Plasmid-borne resistance was was carried by five species, most frequently in *C. nasorum* (5 loci) (Figure 5I). The mobile resistance genes were *ErmX* (macrolide-lincosamide-streptogramin B) and *tet(W)* (tetracycline). The chromosomal AMR repertoire additionally comprised *cmx* (phenicol), *sul1* (sulfonamide) and the aminoglycoside phosphotransferases *APH(3’’)-Ib, APH(3’)-Ia* and *APH(6)-Id* (Figure 5B). By contrast, virulence factors were depleted on mobile loci. Only 6 of 834 VF loci (0.72%) were mobile, a 6.6-fold under-representation (95% CI 1.6 to 9.2-fold depletion, p=4.1×10^-4^). Mobile VFs were restricted to insertion sequences and not horizontally acquired elements (0 of 834 loci on plasmids, prophages or integrons; p=5.6×10^-9^) (Figure 5A, I). These six loci were spread across three species, *C. accolens* (3), *C. guido* (2) and *C. nasorum* (1), and encoded ABC-transporter (4 loci), nitrate-reductase and SpaD-type pilus genes; four of the six retained a chromosomal counterpart (Figure 5A, I).

## 4. Discussion

This work applies standardised genome alignment boundaries to correct species labels amongst publicly accessible *Corynebacterium* genomes (n=5,055; Figure 2A). In clarifying within genus phylogeny, we assigned a species label to 310 NCBI accessions with only a genus or strain level identity, and corrected species assignments for 266 genomes (Figure 2F). We characterised 87 *Corynebacterium* genomes from the sinuses of 71 Australian patients (CRS n=51; non-CRS n=20), and identified four novel-candidate species, pending type strain deposition. Of the publicly accessible genomes, 39 were attributable to our novel lineages, including *C. guido* (n=24 existing genomes / n=10 cohort genomes), *C. hugonis* (n=10 / n=1), *C. lilia* (n=3 / n=1), and *C. lucere*, (n=2 / n=1) [Figure 1D, 2H]. Although metabolic functions were largely conserved amongst sinonasal corynebacteria, there are clear differences between species-complexes, species and even strains. We characterised this variation in metabolism, virulence, and antimicrobial resistance; encoded capacities that mediate competition and mutualism with neighbouring pathogens and commensals.

Species-level microbiome study resolves a more accurate picture of the interactions that occur between microbes occupying the same niche (Lang and Benbow, 2013; Virgo et al., 2025). However, we find that standard methods (MALDI-TOF MS) were unable to resolve sinus corynebacteria at the species level; in the present study, MALDI-TOF MS mis-identified almost half of the cohort (41 of 87 isolates; Figure 1A). However, MALDI-TOF MS assignments accurately sorted isolates along species-complex boundaries (complex-level concordance 87/87; Figure 1A). Thus, in the absence of whole genomes, existing studies of such corynebacteria likely represent findings at the species-complex level. Of note, 17 isolates that MALDI-TOF MS assigned to *C. tuberculostearicum* were resolved to *C. nasorum* by WGS in the present study (Figure 1A). This calls into question published associations between chronic rhinosinusitis and *C. tuberculostearicum*. Despite frequent description in sinus microbiome literature, none of the 41 public *C. tuberculostearicum* genomes were isolated from the sinonasal cavity (Abreu et al., 2012; De Boeck et al., 2019; Escapa et al., 2018). Of the closely related *C. kefirresidentii*, only 2 of 21 genomes were isolated from the sinonasal cavity. In fact, in this study we contribute the first genome from a *C. tuberculostearicum* sinus isolate. Given its lack of representation, *C. tuberculostearicum* is unlikely to be a ubiquitous member of the sinus microbiome. With our augmented annotation of genomes from the *tuberculostearicum-*complex, we position *C. nasorum* as the more relevant taxon in the sinuses. As a species label, *C. nasorum* remains to be represented in commonly used databases for taxonomic profiling including SILVA and EzBioCloud, eHOMD and Greengenes (DeSantis et al., 2006; Escapa et al., 2018; Glöckner et al., 2017; Yoon et al., 2017).

This reinforces the importance of up-to-date and phylogenetically accurate databases for microbiome study. Our understanding of the microbiome is directly dependent on the accuracy of taxonomy and quality of references in our databases. These gaps matter most for shotgun metagenomic sequencing, where read-based k-mer classification (Kraken, Bracken) propagates errors from mislabelled reference genomes. The effect is steep: at a k-mer length of 31, only ∼21% of k-mers remain conserved between genomes sharing 95% ANI (i.e. species level match), falling to ∼4.5% at 90% ANI (e^−kd^; Ondov et al. 2016). Therefore, the small ANI differences that distinguish these species can drastically alter read assignment. As mentioned previously, we found 266 genomes carrying an incorrect species label. In regard to species representation, our review of publicly accessible *Corynebacterium* genomes demonstrated a relative dearth of commensals. This is reflective of the literature’s pathogen-centric bias, whereby sequencing studies have prioritised well-known and often more easily cultured pathogens. As we attempt to understand the roles of these lesser-sequenced commensals, and detect their presence in the microbiome, it is increasingly important to correct this bias in our databases (Han and Vaishnava, 2023). Overall, our findings emphasise the need for a sinus-specific database suitable for metagenomic surveys, as already exists for the gut, oropharynx, skin and vaginal microbiomes (Dai et al., 2022; Escapa et al., 2018; Huang et al., 2024; Liu et al., 2025).

In the human sinus corynebacteria, encoded functions of central biosynthetic pathways, including those involved in nucleotide, amino-acid and carbohydrate metabolism were near-universally complete. However, selected functions varied along phylogenetic lines. This distinction between a conserved biosynthetic core and a variable repertoire for acquiring bioavailable nutrients is consistent with host-niche specialisation (D’Souza et al., 2014). These variable functions can be summarised into themes of (1) carbon, lipid and cofactor acquisition; (2) iron, haem and siderophore economy; (3) nitrogen, nitrate respiration and nitric oxide; and (4) surface factors, stress defence and mobile elements.

The *tuberculostearicum*-complex was distinguished by its expanded capacity for glycan storage. Consistent with this encoded capacity, glycogen accumulation in these species has been demonstrated in vitro (Tran et al., 2024). The closely related UDP-ManNAcA pathway (M01014; wecB/wecC) was almost uniquely encoded by this *tuberculostearicum*-complex, and has been associated with biofilm formation in other respiratory taxa, warranting further investigation (Jurcisek et al., 2005; Tremblay et al., 2013). Host-glycan foraging using the sialic acid operon (*nanAEK*) was carried by all 11 *pseudodiphtheriticum*-complex genomes and by 8 of 31 *C. accolens* isolates. In these sinus commensals, catabolism of free Neu5Ac in mucin via *nanAEK* may limit the nutrient pool available to pathogens such as *S. aureus*, *S. pneumoniae* and *Haemophilus* influenzae (Hatcher et al., 2016; Johnston et al., 2010; Olson et al., 2013). Unlike the *pseudodiphtheriticum*-complex, the *accolens*- and *tuberculostearicum*-species complexes lacked encoded capacity for fatty acid biosynthesis (M00083), necessitating these lipid auxotrophs to scavenge for free fatty acids in the host niche via production of secretory lipases. The free fatty acids produced secondary to lipase secretion have been shown to inhibit the growth of *Streptococcus pneumoniae* (Bomar et al., 2016). While Bomar et al. showed *C. accolens* does this using the secretory lipase LipS1, we found LipS1 to be universally absent in the *accolens*-complex, and broadly present across *pseudodiphtheriticum*-complex and *tuberculostearicum*-complex genomes. Instead, *C. accolens* encoded *estA1*, a GDSL-family lipase/acylhydrolase which could plausibly perform the same role.

Central carbon metabolism also differed along phylogenetic lines. The *accolens*- and *tuberculostearicum*-complexes encoded the acetate-excreting phosphotransacetylase acetate kinase pathway (M00579) whilst the *pseudodiphtheriticum*-complex encoded the glyoxylate cycle (M00012) which permits assimilation of two-carbon compounds (such as acetate) as a carbon source. This raises the possibility of acetate cross feeding, whereby overflow acetate from one lineage serves as a carbon source for another. Metabolite cross-feeding between nasal corynebacteria and *Dolosigranulum pigrum* is well documented and our findings extend this to within-genus mutualism (Brugger et al., 2020; Flores Ramos et al., 2021). This interpretation is further supported by the frequent recovery of multiple *Corynebacterium* species from the same patient (Tran et al., 2024). In terms of interaction with the host immune system, increased short chain free fatty acids in the respiratory tract have been associated with decreased type-2 eosinophil-driven inflammation, and shown to limit eosinophil migration and survival in vitro (Tanabe et al., 2025; Theiler et al., 2019). In the gut, acetate promotes tight junction protein expression in host epithelium, strengthening the mucosal barrier (Kim et al., 2025).

B-vitamin cofactor biosynthesis was the most variable module class, consistent with reliance on exogenous, community-supplied cofactors rather than self-sufficient synthesis. In the *tuberculostearicum*-complex, de novo biotin biosynthesis (M00577) was encoded in only 10 of 17 *C. nasorum* genomes. Loss of this pathway is a recurrent feature of Actinomycetota, many of which lack de novo biotin biosynthesis, instead retaining high affinity biotin uptake systems (Magnúsdóttir et al., 2015; Rodionov et al., 2019). The nasal environment is particularly biotin sparse, making these auxotrophs dependent on biotin released by other nasal commensals (Bilici et al., 2025). This genomic signature is partially corroborated by Bilici et al (2025), who found incomplete biotin-biosynthesis operons in both *accolens-* and *tuberculostearicum*-complex nasal isolates (Bilici et al., 2025). These species thus compete for biotin with *Staphylococcus aureus*, whose own capacity to dominate this niche is equally dependent on its capacity for biotin biosynthesis and high-affinity acquisition (Bilici et al., 2025). The *pseudodiphtheriticum*-complex additionally lacked the thiamine salvage module (M00899), a further axis of cofactor auxotrophy. Collectively, these dependencies position nasal C*orynebacteria* as commensals that have adapted to persist within an interdependent community (D’Souza et al., 2014).

Iron acquisition is a key axis of inter-species competition in the iron-limited nasal niche and our isolates encoded various mechanisms. Every isolate carried *fhuC*, the same ATPase used by *S. aureus* to increase hydroxamate- and staphyloferrin-uptake. This siderophore scavenging encoded in our isolates could compromise *S. aureus’* own fitness under iron limitation, offering a mechanistic basis for Corynebacterium-mediated antagonism of *S. aureus* growth, and a genomic basis for the consumer/cheater interactions demonstrated by Zhao et al. (2024). Beyond iron acquisition, biosynthesis of non-alpha poly-amino acids (NAPAA) metabolites were broadly conserved (76/87 isolates, absent from the *pseudodiphtheriticum-*complex), facilitating the production of polymers such as ε-poly-L-lysine, well recognised for their antibacterial activity, offering a further avenue of *Corynebacterium* competition (Cunningham et al., 2025).

Of all human microenvironments, the paranasal sinuses are particularly rich in nitric oxide (Bazak et al., 2020). Whilst species of the *accolens-* and *pseudodiphtheriticum-* complex encoded a complete nitrate reductase operon (*narGHJI*), this function of anaerobic nitrate respiration was largely absent from *tuberculostearicum*-complex genomes. The process of nitrate reduction yields nitrite and nitric oxide (NO) as intermediates, yet the primary NO detoxifier, flavohaemoglobin (*hmp*), and its regulator (*nsrR*) were restricted to seven genomes (*C. pseudodiphtheriticum, C. bovis,* and *C. lucere*). Most nitrate-respiring sinonasal genomes therefore lack a characterised nitric oxide detoxification system, which may confer nitric oxide sensitivity in these lineages.

Across the cohort, virulence and antimicrobial resistance was minimally encoded, and all species lacked the diphtheria toxin (*tox*) gene. The two virulence factors that were shared across all isolates were housekeeping proteins, more indicative of conserved core machinery than pathogenic capacity. Against this shared baseline, differences in AMR genes and virulence were predominantly observed at the species level, with 17 of 22 rare virulence factors confined to single species. Among well sampled species, *C. guido, C. nasorum* and *C. rhinophilum* had the lowest virulence density of in their genomes, whilst *C. pseudodiphtheriticum* carried the greatest density of both virulence and resistance determinants. Study of mobile-element content demonstrated horizontal gene transfer of AMR genes but not virulence. Resistance genes were disproportionately mobile: roughly one in six sat on a plasmid, more than thirty time the expected share of coding sequences, and almost none had a chromosomal copy to fall back on. Prevalence of mobile compartments were strongly species and even strain dependent. Insertion-sequence load ranged from 15.5 in *C. pseudodiphtheriticum* to 127.5 in *C. rhinophilum*. The pronounced insertion-sequence expansion in *C. rhinophilum* as well as other species from the *accolens*-complex, is consistent with the intra-genome proliferation and genome capacity reduction. Rather than horizontal gene acquisition, insertion sequences represent genomes redundancy, and their proliferation is seen in bacteria as they adopt increasingly host-restricted lifestyles, an early signature of niche-adapted commensal evolution (Siguier et al., 2014).

A key limitation of this study is the speculative nature of the identified genomic differences in the absence of confirmatory in vitro testing. As such, we propose our findings in differential gene presence as starting points for further interrogation. In particular, confirmatory in vitro study of the proposed differences in metabolism between these species as well as their biofilm forming capacity, cytotoxicity, and examination of antibiotic resistance profiles would substantiate the genome-based findings of this article. Additionally, our four novel candidate species (*C. guido* [nearest accolens], *C. hugonis* [nearest rhinophilum], *C. lilia* [no taxon within 90% ANI] and *C. lucere* [nearest amycolatum]), also require further study to characterise morphology, growth and metabolism and submission of type strains.

## 5. Conclusion

In conclusion, this study addresses the underexplored diversity within the *Corynebacterium* genus. Functionally, corynebacteria diverged along phylogenetic lines in accessory metabolism, particularly in relation to glycogen and glycan utilisation, fatty-acid biosynthesis, cofactor auxotrophy and iron acquisition. At the species level, sinus corynebacteria encoded different survival strategies supporting mutualism within the genus and competition against pathogens such as *S. aureus* and *S. pneumoniae*. These sinus isolates showed low virulence and AMR carriage, with virulence factors conserved down the *Corynebacterium* lineage rather than horizontally acquired. By re-deriving species boundaries across the entire genus, we provide a type-strain anchored taxonomy that reassigns hundreds of mislabelled or unresolved public accessions. Furthermore, we resolve the mischaracterisation and conflation of species within the *tuberculostearicum*-complex that has permeated the sinus microbiome literature. We contribute 87 new genomes, including four candidate novel species. Together, these findings point to common *Corynebacterium* as sinonasal commensals, a role now open to confirmation by species-resolved microbiome study.

## Data Summary

1. GenBank bio project
2. Type strain deposition
3. Code availability

## Acknowledgements

NA

## CRediT author contribution statement

S.V., P.J.W. and A.J.P. conceived and designed the study. L.D., C.C., E.B., N.C.V. and J.P.P. assisted with laboratory work. I.A.B. conceived and designed the study, performed the laboratory work, developed the computational pipeline, analysed the data, produced the figures and wrote the manuscript. All authors reviewed and approved the manuscript.

## Declaration of generative AI and AI-assisted technologies in the manuscript preparation process

We acknowledge the use of generative AI in this article for the purpose of checking spelling, grammar and references (to manuscript data and external sources); also to refine and compile scripts written by the author into python notebooks to accompany this article for code availability, these notebooks were cross checked independently by the author.

## Data availability statement

Pipeline code: https://gitlab.com/isabella-burdon/taxanalyser.git

Statistics code: https://gitlab.com/isabella-burdon/corynebacterium_study.git

Whole genome data: BioProject ID pending

## Supplementary materials

**Supplementary Figure 1.**
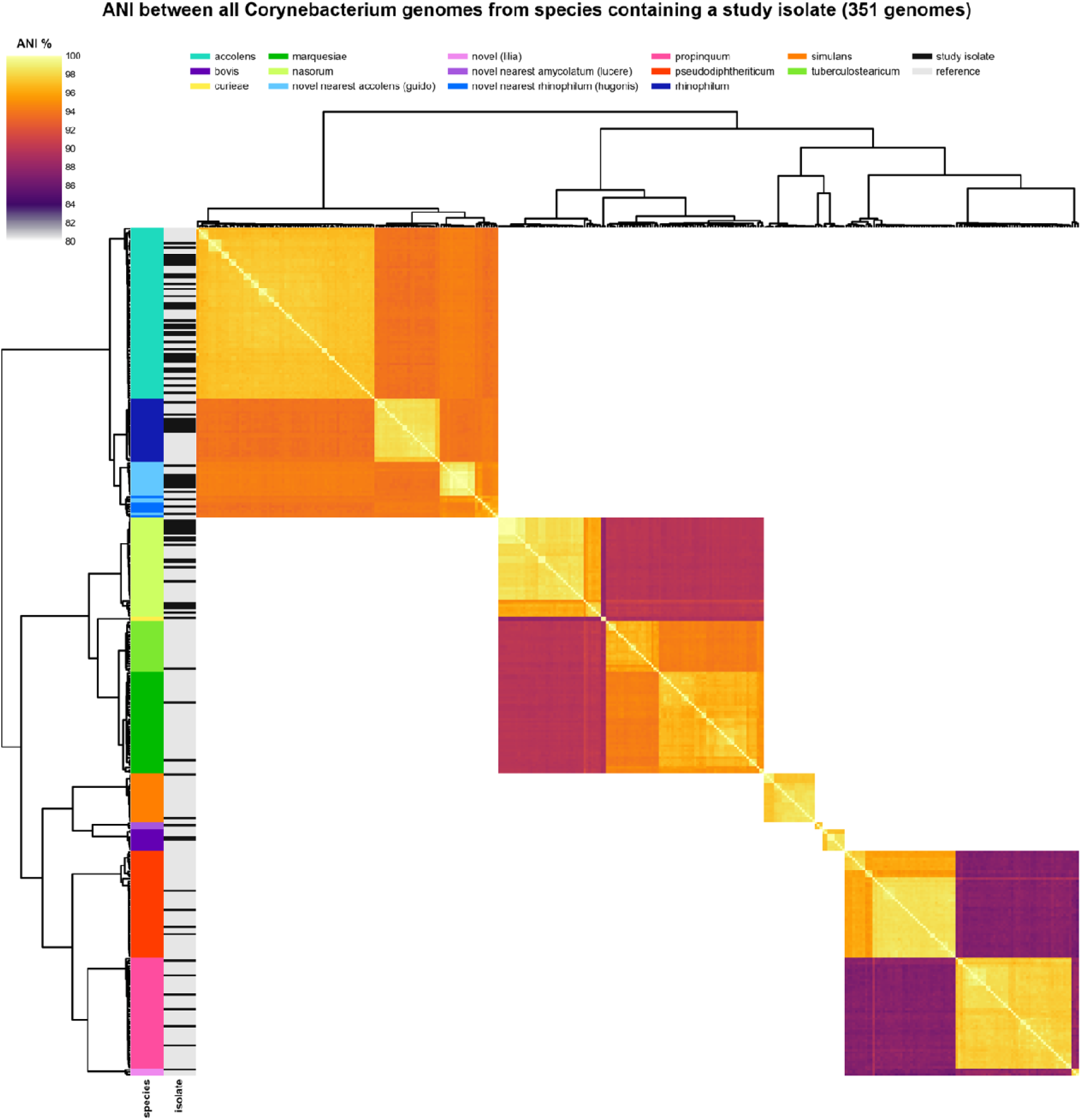
All-versus-all ANI matrix of quality filtered Corynebacterium genomes (NCBI accessions + clinical isolates) showing species that were represented by an isolate in the current sinus Corynebacterium cohort.

**Supplementary Figure 2.**
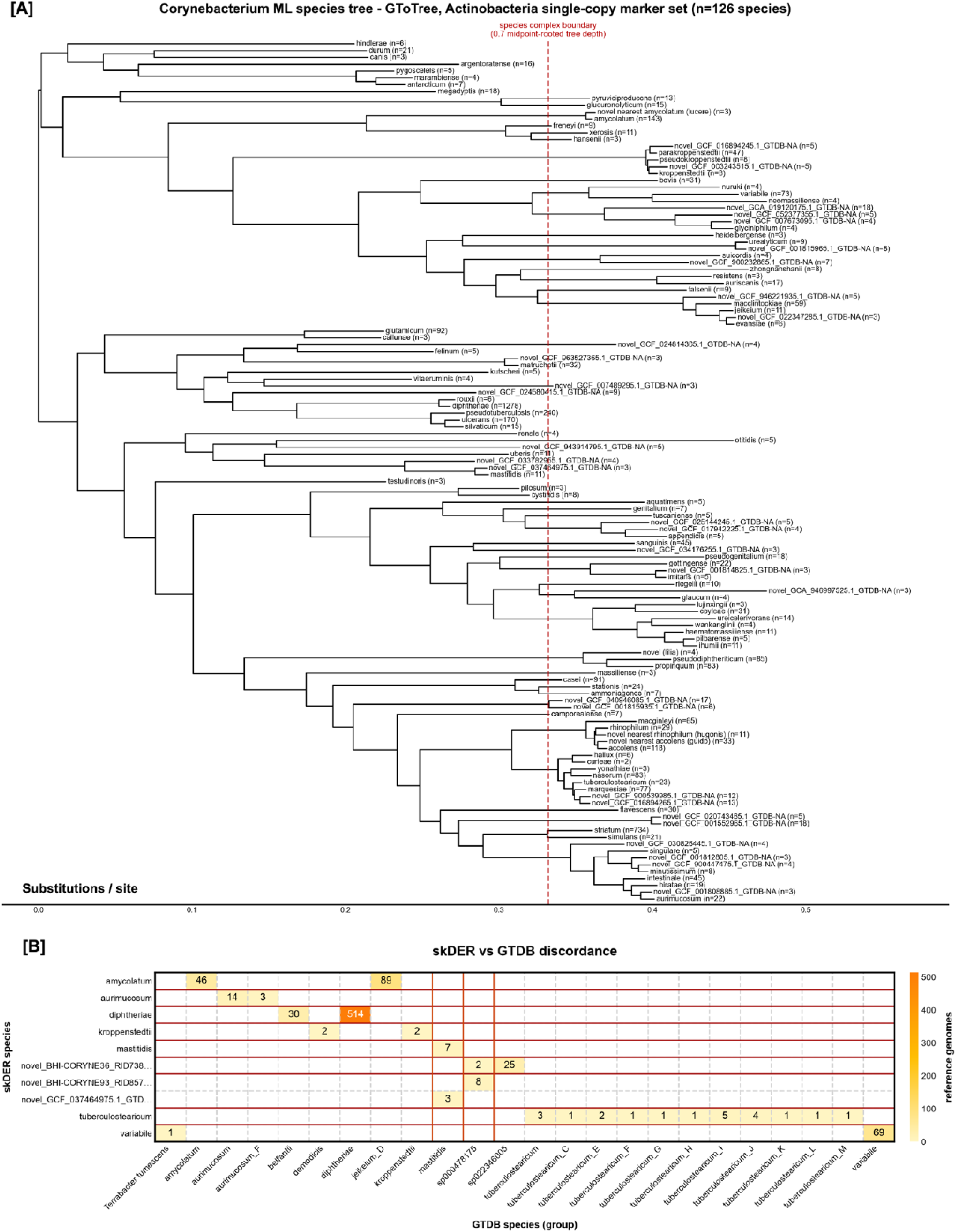
(a) Midpoint-rooted maximum-likelihood phylogeny of the Corynebacterium genus (128 species) inferred with GToTree from the Actinobacteria single-copy marker-gene set. Each tip is a species defined by the de novo (skDER) respeciation, with the number of genomes per species in parentheses; the red dashed line marks the species-complex boundary at 70% of the midpoint-rooted tree depth. (b) Discordance between skDER species assignments (rows) and GTDB genomospecies (columns), shown as the number of genomes assigned to each combination; only skDER species with at least one discordant assignment are shown. Most discordance reflects GTDB placeholder genomospecies names (e.g. tuberculostearicum_C–M) rather than disagreement over species boundaries.

**Supplementary Figure 3.**
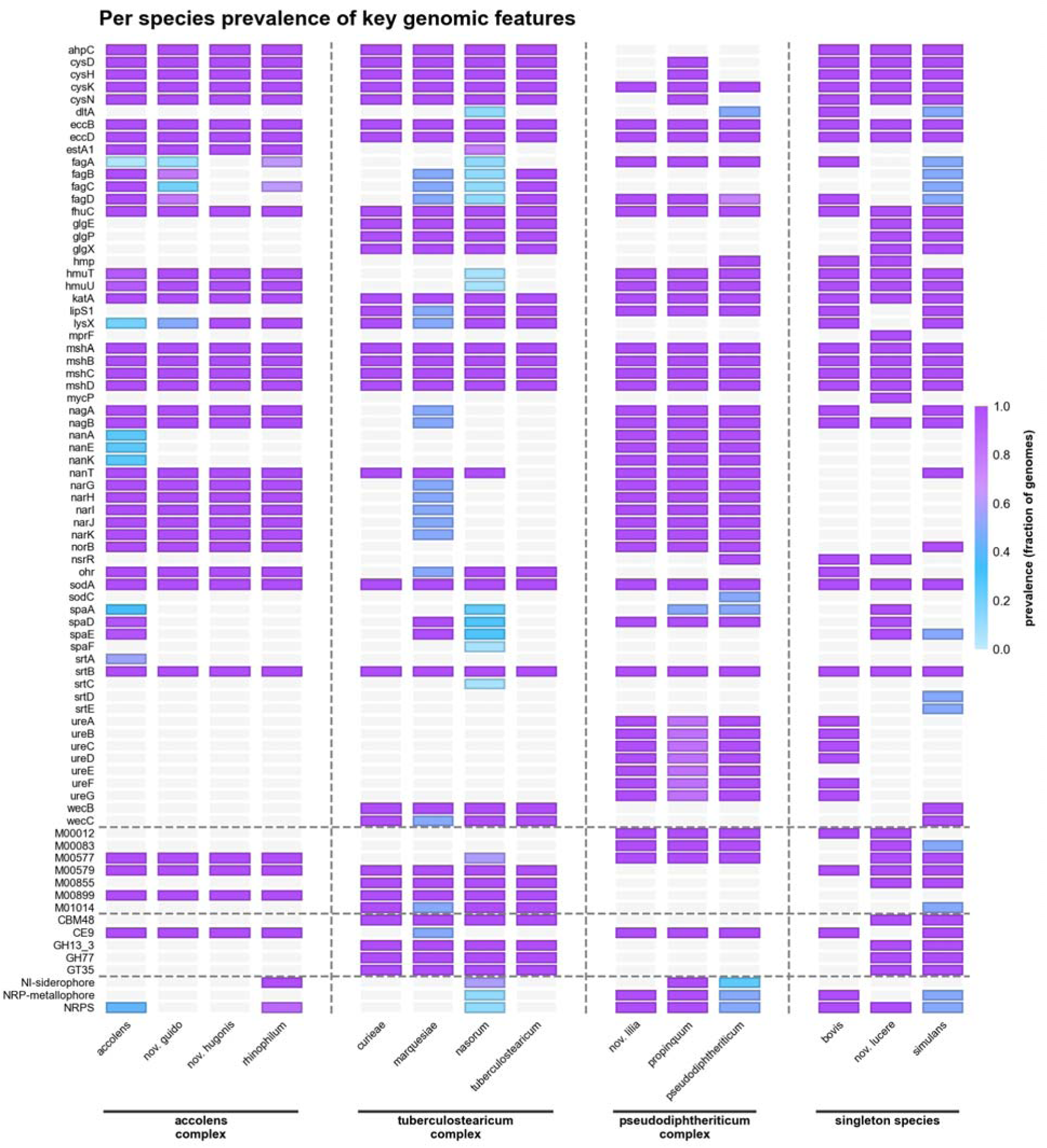
Prevalence of the reported genomic features, shown per species and grouped by species complex. Heatmap of the prevalence (fraction of genomes; colour scale) of the features reported in the Results and Discussion, grouped by feature type and separated by dashed rules: 63 genes, 7 KEGG modules, 5 CAZy families and 3 biosynthetic gene cluster classes, alphabetically within type; pale grey marks prevalence 0. Gene symbols are the annotation consensus, with the eggNOG preferred name where the consensus leaves a CDS unnamed; a module is counted complete at ≥0.8 block completeness, and a CAZy family or BGC class present at a count above zero. This panel is descriptive; no statistical test was applied.

**Supplementary Figure 4.**
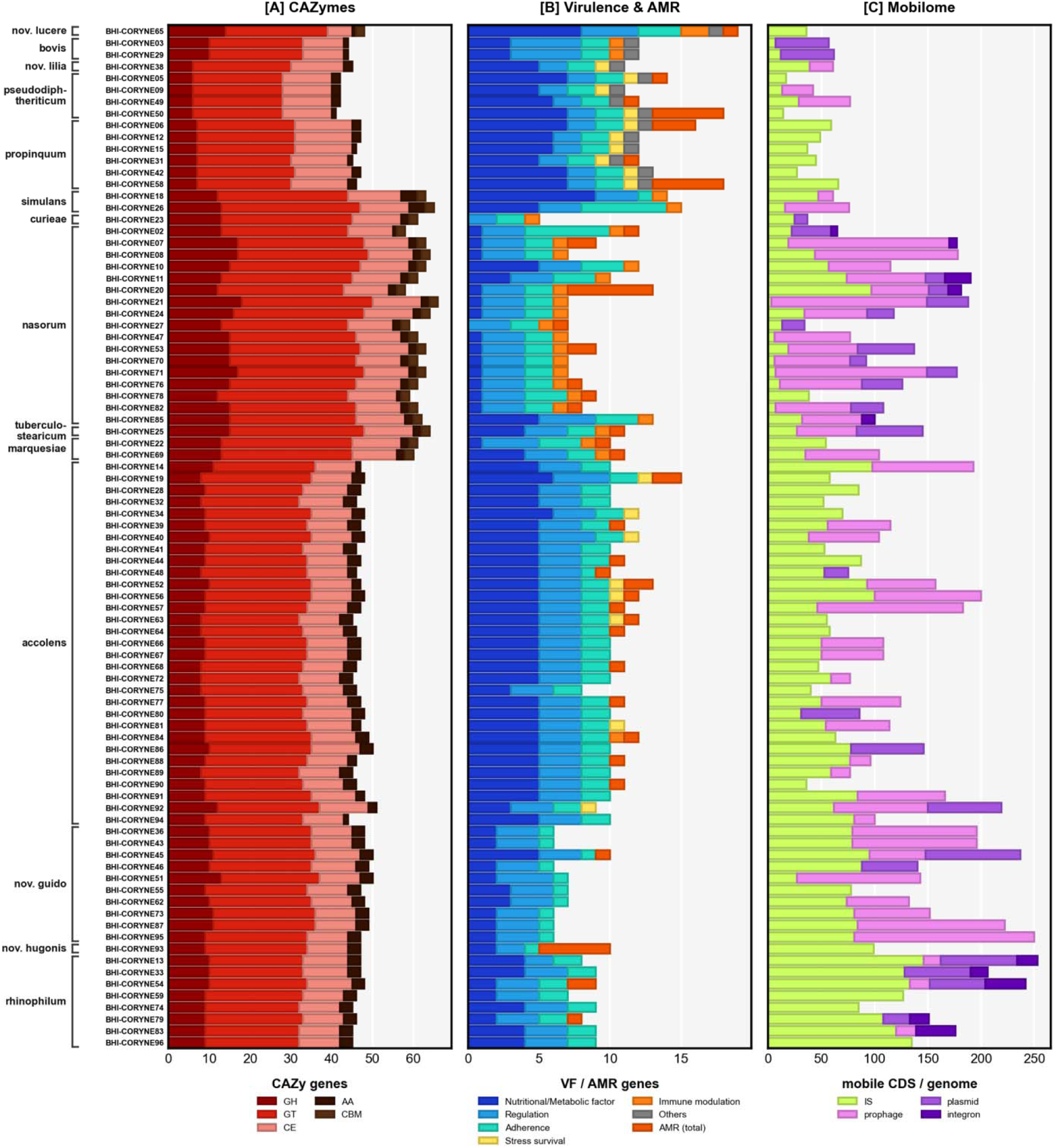
Per-genome counts of functional and mobile-genetic-element content across the 87 sinus Corynebacterium isolates, grouped by species (labelled at left). (a) Carbohydrate-active enzyme (CAZyme) genes, coloured by class: glycoside hydrolases (GH), glycosyltransferases (GT), carbohydrate esterases (CE), auxiliary activities (AA) and carbohydrate-binding modules (CBM). (b) Virulence factor (VF) and antimicrobial-resistance (AMR) genes, coloured by functional category (nutritional/metabolic factor, regulation, adherence, stress survival, immune modulation, other) with total AMR genes shown separately. (c) Mobile-genetic-element content (mobile coding sequences per genome): insertion sequences (IS), prophages, plasmids and integrons. Each horizontal bar is one isolate.

